# Fractone bulbs derive from ependymal cells and their laminin composition influence the stem cell niche in the subventricular zone

**DOI:** 10.1101/093351

**Authors:** Marcos Assis Nascimento, Lydia Sorokin, Tatiana Coelho-Sampaio

**Author notes:** **Corresponding author** Marcos Assis Nascimento, Address: Av. Carlos Chagas Filho 373, Centro de Ciências da Saúde, B-11, Rio de Janeiro, Brazil. Postal code: 21941-902.

## Abstract

Fractones are extracellular matrix structures in the neural stem cell niche of the subventricular zone (**SVZ**), where they appear as round deposits named bulbs or thin branching lines called stems. Their cellular origin and what determines their localization at this site is poorly studied and it remains unclear whether they influence neural stem and progenitor cells formation, proliferation and/or maintenance. To address these questions, we analyzed whole mount preparations of the lateral ventricle by confocal microscopy using different extracellular matrix and cell markers. We found that bulbs are rarely connected to stems and that they contain laminin α5 and α2 chains, respectively. Fractone bulbs were profusely distributed throughout the SVZ and appeared associated with the center of pinwheels, a critical site for adult neurogenesis. We demonstrate that bulbs appear at the apical membrane of ependymal cells at the end of the first week after birth. The use of transgenic mice lacking laminin α5 gene expression (*Lama5*) in endothelium and in FoxJ1-expressing ependymal cells, revealed ependymal cells as the source of laminin α5-containing fractone bulbs. Loss of laminin α5 from bulbs correlated with a 60% increase in cell proliferation, as determined by PH3 staining, and with a selective reduction in the number of quiescent neural stem cells in the SVZ. These results indicate that fractones are a key component of the SVZ and suggest that laminin α5 modulates the physiology of the neural stem cell niche.

**Significance Statement:** Our work unveils key aspects of fractones, extracellular matrix structures present in the SVZ that still lack a comprehensive characterization. We show that fractones extensively interact with neural stem cells, whereas some of them are located precisely at pinwheel centers, which are hotspots for adult neurogenesis. Our results also demonstrate that fractones increase in size during aging and that their interactions with NSPCs become more complex in old mice. Lastly, we show that fractone bulbs are produced by ependymal cells and that their laminin content regulates neural stem cells.

## Introduction

The subventricular zone (**SVZ**) is the largest and most studied neural stem cell niche in adult mice. Within this niche, neural stem and progenitor cells (**NSPCs**) contact the ependymal cell layer lining the ventricles (Doetsch et al., 1999) and the basement membranes (**BMs**) of blood vessels (Shen et al., 2008; Tavazoie et al., 2008). BMs are specialized extracellular matrix sheets, which in most tissues underlie endothelia and epithelial layers and ensheath nerves and muscle fibers, but in the SVZ appear in an unique fractal-like pattern. In the CNS, blood vessels are surrounded by two biochemically distinct BMs, one underlying the endothelium and one produced by astrocytes and deposited in the parenchymal BM (Sixt et al., 2001; Wu et al., 2009). Fractones are described to emerge from capillaries as thin stems, which branch profusely and terminate as BMs bulbs at the ependymal layer (Leonhardt and Desaga, 1975; Mercier et al., 2002, 2011), thereby accounting for their punctate appearance. Laminins are the main functional constituents of BMs, acting as a molecular scaffold for its assembly and providing biological signals that control cell migration, differentiation and proliferation (Smyth et al., 1999). All members of the laminin family are heterotrimers composed of one α, one β and one γ chain that combine to form 16 distinct isoforms with tissue specific distributions and functions (Durbeej, 2010). The C-terminus of the 5 laminin α chains are key to laminin signaling, bearing in most isoforms the domains recognized by membrane receptors that regulate cellular adhesion, migration and division (Colognato and Yurchenco, 2000). Laminins containing the α5 chain have been shown to be crucial for pluripotent stem cell survival and self-renewal *in vitro* (Hongisto et al., 2012; Miyazaki et al., 2012; Laperle et al., 2015) and for inhibiting proliferation *in vivo* (Wegner et al., 2016). The effects of fractones on NSPCs physiology has been attributed to the perlecan; however, this is based solely on its ability to bind growth factors, e.g. bFGF, BMP-4 and BMP-7, and to present them to contacting cells (Douet et al., 2012; Kerever et al., 2014; Mercier and Douet, 2014).

Despite growing evidence of the relevance of fractones in the SVZ (Kazanis et al., 2010; Mercier et al., 2012), a comprehensive understanding of their location is still lacking, in particular how fractones relate to NSPCs and pinwheels, critical structures for adult neurogenesis formed by NSPCs and ependymal cells at the ventricle wall (Mirzadeh et al., 2008). In addition, identifying the cells responsible for producing fractones and determining whether the laminin composition plays a role in SVZ physiology are crucial to understanding how stem cells are formed and maintained at this site.

Previous studies have identified interactions between GFAP^+^ neural stem cells and pan-laminin+ fractones using electron microscopy and immunofluorescence stainings of coronal sections of the SVZ (Leonhardt and Desaga, 1975; Mercier et al., 2002, 2011). Although suitable for unveiling the existence of such interactions, analyzes of thin tissue sections cannot fully address interactions between fractones and neural stem cells at the SVZ, due the filamentous nature of GFAP+ cells. We therefore here analyze the position of fractone bulbs using 3D reconstruction of immunofluorescently stained whole mounts of the lateral wall, revealing that bulbs are frequently located precisely at the center of pinwheels. To identify the cell responsible for producing fractone bulbs and to investigate whether its laminin composition is important for regulating NSPC proliferation in the SVZ, we analyzed mice lacking *Lama5* expression in endothelial cells or ciliated ependymal cells. Our data indicate that ependymal cells are the source of laminin α5-containing fractones, and demonstrated that the loss of laminin α5 at this site correlated with a 60% increase in overall proliferation of NSPCs and to a decrease in the number of quiescent neural stem cells. Our findings indicate that fractone bulbs are ependymally-derived basement membrane structures critical to SVZ physiology.

## Materials & Methods

### Experimental Animals

FoxJ1-Cre::*Lama5*^−/−^ mice were obtained by crossing FoxJ1-Cre mice (Zhang et al., 2007) with *Lama5*^flox/flox^ mice (Song et al., 2013); Tek-Cre::*Lama5*^−/−^ mice were obtained in analogous way (Song et al., 2013). All mice were on a C57BL/6 background (Charles River Laboratories).Male and female mice were used. Animal breeding and procedures were conducted according to the German Animal Welfare guidelines.

### Histological Preparations

Brains were dissected and fixed overnight at 4°C in 2% paraformaldehyde in PBS. Thick (50 µm) coronal sections were made using a Zeiss Vibratome. Whole mounts were prepared from dissecting lateral walls of the lateral ventricles as described previously (Mirzadeh et al., 2008, 2010).

### Antibodies & Immunofluorescence

Sections and whole mounts were washed in PBS, blocked with 5% goat serum/PBS or 1% BSA/PBS (Sigma) and incubated with primary antibodies overnight at 4°C. Primary antibodies used: Anti-laminin α2 (4H8-2) (Schuler and Sorokin, 1995) anti-laminin α5 (4G6) (Sorokin et al., 1997b), anti-laminin γ1 (3E10) (Sixt et al., 2001), anti-pan-laminin(455) (Sorokin,1990), anti-β-catenin (Sigma cat.# C2206, 1:1000), anti-GFAP (Sigma cat.# C9205, 1:800; eBioscience cat.# 53-9892, 1:500), anti-nestin (rat-401, DSHB/Iowa 1:100), anti-PH3 (Millipore cat.# 06-570, 1:500). After the incubation, tissues were thoroughly washed with PBS and incubated with secondary antibodies: Alexa 488 anti-rat (Life Technologies cat.# A11006, 1:1000), Alexa 555 anti-rat (Abcam cat.# ab150154, 1:1000), Alexa 568 anti-rabbit (Life Technologies cat.# A11011 1:1000), Alexa 647 anti-rabbit (Jackson/Dianova cat.# 111-605-144 1:1000). After incubation with secondary antibodies, thick sections and whole mounts were mounted with mounting media and a coverslip. Tissues were analyzed using a Zeiss AxioImager microscope equipped with epifluorescent optics and documented with a Hamamatsu ORCA ER camera or with a Zeiss confocal laser scanning microscope LSM 700.

### In situ hybridization

In situ hybridization for laminin α5 mRNA was performed as previously described (Sorokin et al., 1997a).

### Pulse-chase assay for Edu

3 months old mice received intra-peritoneal injections of 50 mg/kg EdU (Click-iT ^®^ EdU Alexa Fluor 647 Imaging kit; Thermo catalog # c10340) in sterile PBS at three consecutive days. Whole mounts of the ventricular lateral walls were prepared after six weeks, scanned in a Zeiss Axio Imager M2 with an automated stage and EdU-labeled cells were counted using ImageJ software.

### Morphological and Statistical Analyses

For PH3+ nuclei quantification, whole mounts were photographed with a 10x objective lens in the Zeiss AxioImager. Pictures were merged using Adobe Photoshop CS3 (Adobe Systems, Mountain View, CA). Merged pictures were processed using ImageJ (U.S. National Institutes of Health, Bethesda, MD, USA) and nuclei were quantified. For quantification of fractone bulbs volume, pan-laminin stainings in coronal sections were rendered in tridimensional volumes and quantified using Imaris 7.0 software (Bitplane). For analyses of interactions of GFAP+ processes and bulbs in old mice, GFAP and pan-laminin stainings of whole mounts were reconstructed in 3D using Imaris. Statistical analyses were made using Graphpad Prism 6 (GraphPad Software Inc., La Jolla, CA). A t test with a 95% confidence interval was used to analyze statistical differences in figures 4B, 5B, 6B-C.

## Results

### An en face view of interactions between GFAP+ cells and fractones

In order to have a more comprehensive view of the interactions between neural stem cells (NSCs) and fractones, whole mounts of the SVZ were simultaneously immunofluorescently stained for laminin ²1 to mark BMs, GFAP to mark NSCs and β-catenin to delineate cell contacts at the ependymal layer. First, we observedthat fractones bulbs account for a significant part of all BM in the neurogenic niche (Fig. 1*A*) and are located preferentially at the interface between neighboring cells in the ependymal layer (Fig. 1*B*). GFAP+ neural stem cells interact with BMs of blood vessels (Fig. 1*C*, *yellow arrow*) but their cell bodies and processes also contact several fractone bulbs at the same time (Fig. 1*C*, *arrowheads*).

**Figure 1:**
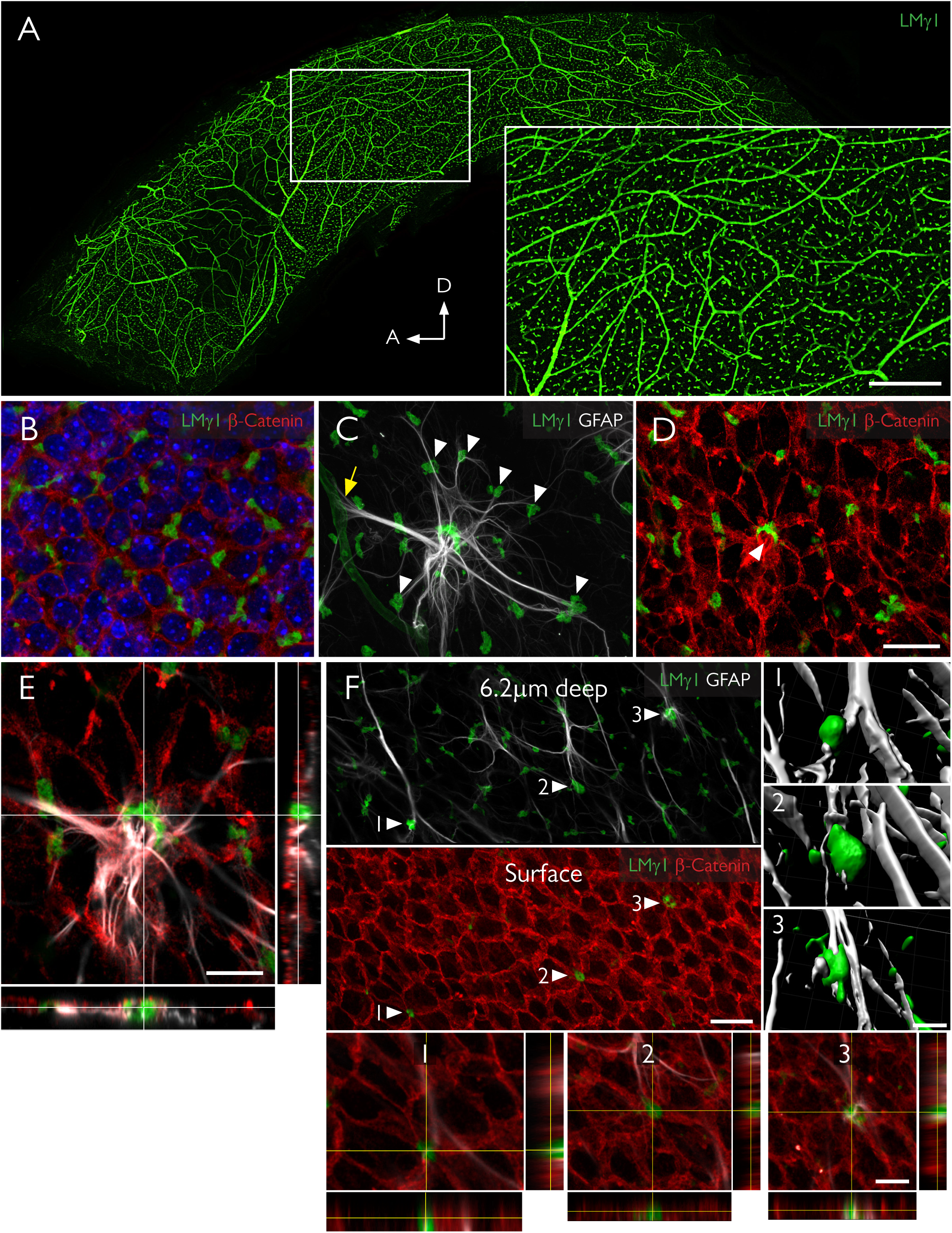
Fractones are a major BM source in the SVZ and appear at the center of pinwheels. ***A***, Whole mount of the lateral ventricle wall of an adult mouse immunofluorescently stained for laminin γ1 (green). The frontal view reveals the profuse distribution of fractone bulbs at the neurogenic niche. A higher magnification of the boxed area is depicted in the inset. ***B***, Frontal view of the ependymal cell layer, seen at 1 µm depth relative to the ventricular surface, stained for laminin γ1 (green) to visualize fractones and β-catenin (red) to visualize cell junctions in the ependymal layer. All fractone bulbs sit at the interface between cells. ***C***, Maximum intensity projection of a Z-stack of a whole mount showing a GFAP+ NSC (gray) in contact with one blood vessel (yellow arrow) and more than 10 fractone bulbs (arrowheads) at the same time. ***D,*** The neural stem cell body also contacts a large bulb located at the center of a pinwheel (arrowhead). ***E***, Orthogonal views revealing that the NSC-fractone interaction seen in C-D occurs at the surface of a pinwheel center. ***F***, GFAP+ processes contact more bulbs at pinwheel centers (arrowheads) than bulbs at the basolateral surface of ependymal cells. Interactions between GFAP+ processes and fractone bulbs were labeled 1–3 and rendered, revealing that GFAP+ processes establish direct and complex contacts with fractones. Orthogonal views of these interactions confirm that fractone bulbs sit between ependymal cells at pinwheel centers and GFAP+ processes directly contact them Scale bars: A: 500 µm, inset: 200 µm. B-D: 25 µm. E: 12 µm. F: 25 µm, orthogonal views: 12µm.

NSCs exhibiting processes that contact the cerebrospinal fluid are the basic neurogenic unit in the SVZ and, together with surrounding ependymal cells, form a distinct pinwheel-like arrangement (Mirzadeh et al., 2008). We observed that fractone bulbs are frequently located at the center of pinwheels, coinciding with GFAP+ hubs from which abundant processes radiate out to make contact with surrounding individual bulbs (Fig. 1*C-E*).These bulbs are aligned with the apical pole of ependymal cells at the ventricular surface (Fig. 1*F*, surface, arrowheads; *G*). Most fractone bulbs occur at the basolateral side of the ependymal cells, where they contact GFAP+ processes (Fig. 1*F*, 6.2 µm deep, arrowheads). Three-dimensional reconstruction and orthogonal views of contact points between GFAP+ processes and bulbs at pinwheel centers (arrowheads labeled 1–3) revealed direct contact between these two structures.

### Fractone bulbs are enriched in laminin α5

Fractones are regarded as BM structures that emerge from capillaries as thin threads, known as stems, that branch out to reach the ependymal layer where they appear as spherical deposits known as bulbs (Mercier et al., 2002, 2011). We used coronal sections of the SVZ and monoclonal antibodies specific for laminin α chains to investigate the laminin isoforms present in fractones. We found that fractone bulbs exhibited strong laminin α5 staining, and only very few contained the α2 chain (Fig.2*A-E*). Laminin α4 was not found in any fractone bulb (Fig.2*A*), in contrast to previous reports (Kazanis et al., 2010). Laminin α1 and α3 chains were also completely absent in bulbs (not shown), which is consistent with the literature (Kazanis et al., 2010). Conversely, the predominant laminin α chain present in stems was laminin α2 (Fig. 2*C; E*) which, in the adult CNS, is produced mostly by astrocytes of the blood-brain barrier (Sixt et al., 2001) and potentially by pericytes (Armulik et al., 2010). Some staining for laminin α4 and α5 was found in stems (not shown). Surprisingly, we found that although a few bulbs contacted stems (Fig. 2*C*; *E*, arrowheads) the vast majority did not touch stems, which together with the observation that they contain different laminin chains, suggests that bulbs and stems are independent structures.

**Figure 2:**
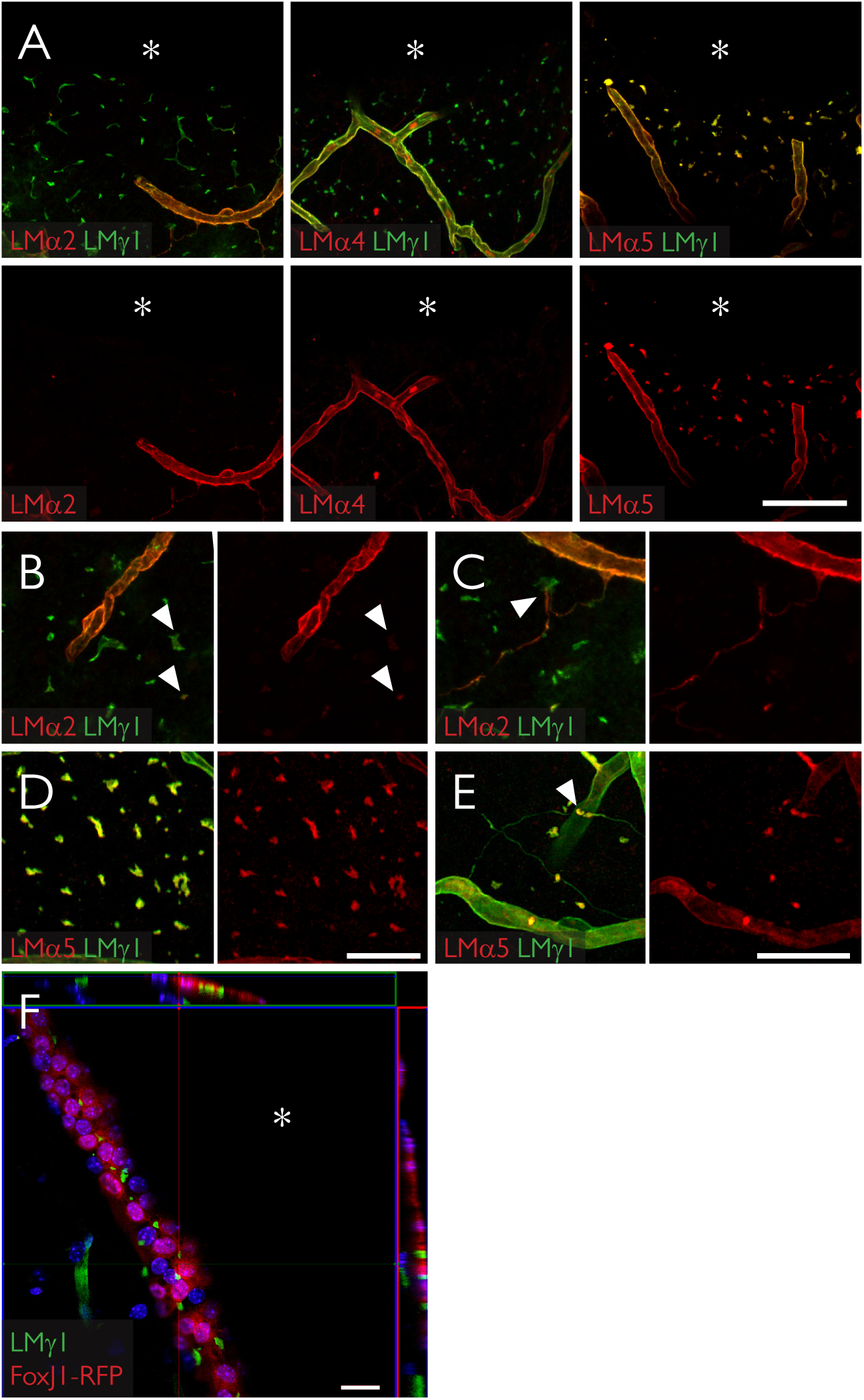
Fractone bulbs and fractone stems have distinct laminin compositions. ***A***, Immunolabeling for laminin α2 (red, left), laminin α4 (red, middle), laminin α5 (red, right) and laminin γ1 (green) seen in maximum intensity projections of thick (60 µm) coronal sections of young adult mice showing that fractone bulbs and blood vessels have distinct laminin compositions. Laminin α5 is the main α chain in fractone bulbs, followed by a faint presence of laminin α2. Laminin α4 is absent from fractones. ***B***, Higher magnification showing the faint presence of laminin α2 in fractone bulbs (arrowheads). ***C***, Higher magnification depicting fractone stems, rich in laminin α2 emerging from a blood vessel and contacting fractone bulbs (arrowhead). ***D***, Higher magnification showing the ubiquitous presence of laminin α5 in fractone bulbs and in ***E*** its absence in most of fractone stems. ***F***, Orthogonal views from a coronal section of a FoxJ1-RFP mouse showing that fractone bulbs are located at the ependymal layer only. Fractones bulbs may appear in thick stripes in coronal sections due the angle of the lateral wall and the thickness of the optical section. Asterisks indicate the ventricular cavity. Scale bars: A: 100 µm; B, D: 50 µm; C, E: 50 µm; F: 20 µm.

To properly interpret images in Figure 2, as well as in following Figures 3 and 5, it is important to emphasize that the wall of the lateral ventricle is not always orthogonal to the coronal axis. As a consequence, the thicknesses of the stripes across which bulbs vary largely between individual sections, despite bulbs being exclusively located between cells in the ependymal layer (Fig. 1*B*). This is verified with FoxJ1-RFP reporter mice that expresses RFP in the ependymal layer (Fig. 2*F*).

### Fractone bulbs are produced by ependymal cells during the first week after birth

To determine when fractone bulbs form and whether this correlates with appearance of laminin α5 expression, we investigated the emergence of fractones at the ependymal walls during development. To do so, we checked the laminin isoform expression at the SVZ every day during the first week after birth. Fractone bulbs, as defined by a punctate laminin γ1 staining, were not detected at P0, however, in addition to blood vessel staining, a faint and disperse laminin γ1 chain staining lining the ventricle was detectable at the nestin+ layer of late radial glial cells/early NSCs (Fig. 3*A*). Laminin α5 expression in blood vessels in the CNS starts approximately at the third week after birth (Sorokin et al., 1997b). However, in situ hybridization showed that mRNA for the laminin α5 chain is already expressed at the ventricle surface (Fig. 3*B,* arrow) and also at the choroid plexus at P0 (Fig. 3*B*, arrowhead) as reported previously (Sorokin et al., 1997b). At P3, more condensed spots positive for laminin γ1 and α5 were present in some regions of the ventricle (Fig. 3*C*). Laminin expression was spatially heterogeneous, with dense laminin expression at regions of the dorsal wall while most portions of the lateral wall lacked laminin immunoreactivity. At P7, early laminin α5 and γ1 chain positive fractones were visible along all the ependymal surface of the lateral ventricles (Fig. 3*D*). In a frontal view of a P7 ependyma using a whole mount preparation, we observed that these first fractones are small patches of BM (Fig. 3*E*) not aligned with regions of cell-cell contact as in adult mice (Fig. 1*B*). Curiously, these early deposits are seen only at the apical surface of the ependymal cells, (Fig. 3*F*). Analysis of FoxJ1-RFP reporter animals that express RFP under the control of the ependymal-specific promoter FoxJ1, revealed laminin γ1 inside ependymal cells and some early fractones already at the cell surface (Fig. 3*G*).

**Figure 3:**
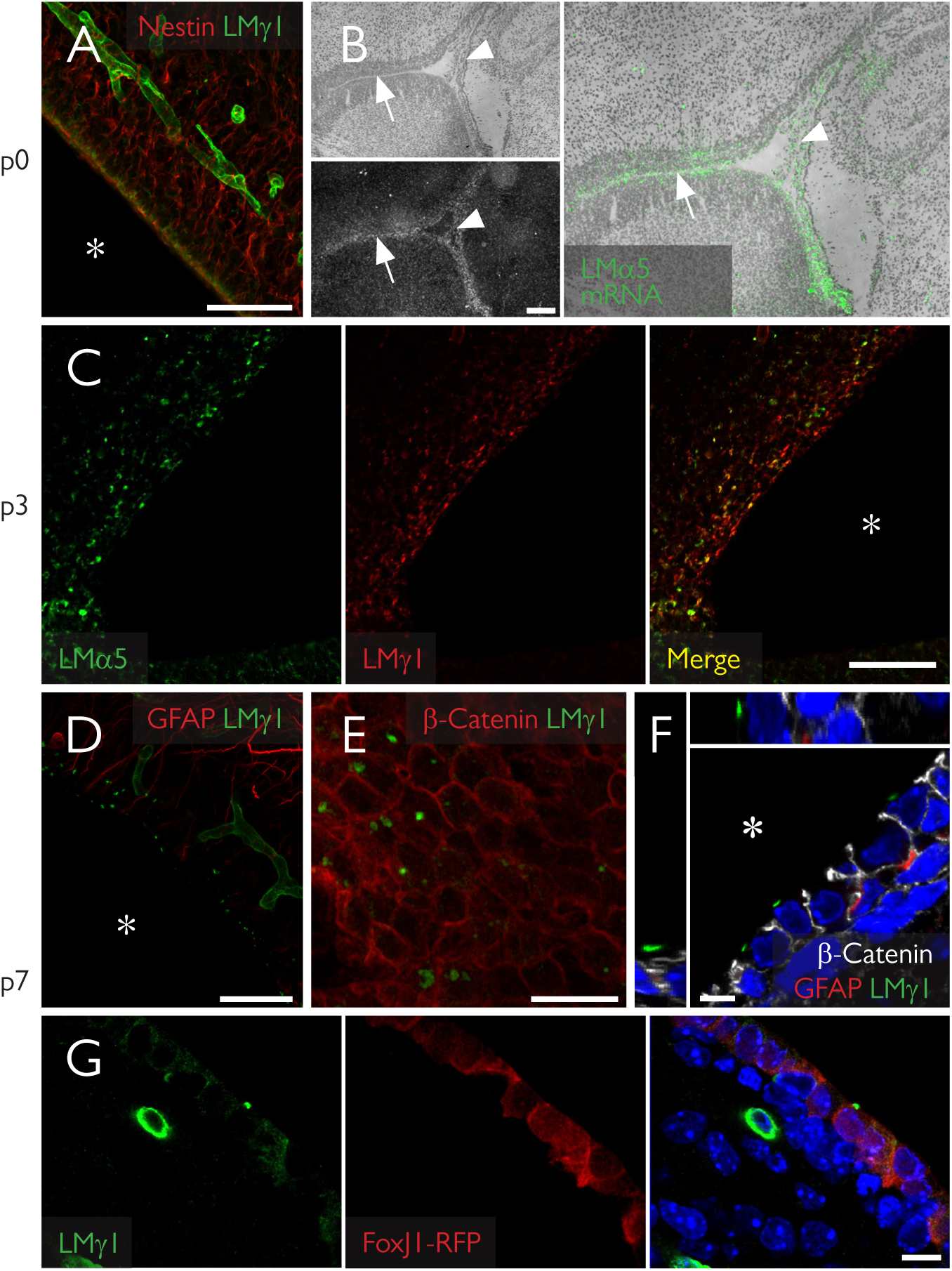
Fractones emerge at the apical surface of ependymal cells in the first week after birth. ***A***, Coronal section of a newborn mouse showing scarce laminin expression (green) at the ependymal layer. ***B***, *In situ* hybridization of a sagittal section at the same age revealing extensive *Lama5* gene expression (gray in bottom panel, green in right panel) at the ependymal layer (arrow) and at the choroid plexus (arrowhead), weeks before protein is detected in blood vessels. ***C***, At day 3, laminin α5 is expressed in a more punctate pattern in several regions of the ependymal surface. ***D***, At day 7, early fractone bulbs can be clearly seen in all the ventricular surface. A frontal view of the ependymal layer (***E***) shows fractones at the apical surface of ependymal cells, confirmed by orthogonal sections of a confocal stack (***F***). ***G***, At the same time, laminin γ1 can be seen both as a punctate pattern at the apical cell surface and a diffuse intracellular stain inside RFP+ ependymal cells in FoxJ1-RFP mice. Asterisks indicate the ventricular cavity. Scale bars: A: 50 µm, B: 200 µm, higher magnification: 130 µm, C: 50 µm, D: 50 µm, E: 25 µm, F: 5 µm, G: 10 µm.

The fact that fractones bulbs first appear as small BM patches at the apical surface of ependymal cells indicates that these structures undergo major spatial and morphological changes until adulthood. We therefore investigated whether fractone size changed over time by measuring their volumes in 3D stacks from confocal microscopy at P34, P86, P162, P175, P269 and P303. We observed that the mean volume increases gradually, showing a linear correlation with age (R²=0.91). This result extends the recent observation that bulbs are larger in 100 week old mice than in 12 week-old mice (Kerever et al., 2015). A frequency distribution analysis showed that this increase in size occurs in the entire fractone population (Fig. 4*A, B*). Mice older than 200 days displayed large fractones bulbs (> 40 µm³) and were associated with more GFAP+ processes contacts than bulbs in younger animals. Old mice (>200 days) displayed fractones with a tunnel-like structure and neural stem cells extend their processes within these tunnels to contact the CSF (Fig. 4*C*). Tunneled bulbs, not seen in younger animals, were often, but not always, located at the center of pinwheels (Fig. 4*C*). In addition, old mice had unique enlarged fractone bulbs, with some reaching over 20 µm in diameter (Fig.4*D*, *arrowheads*). These giant fractones were associated with clusters of cells at the anterior ventral part of the ventricular wall (Fig. 4*D*, *arrows*). Such cell clusters were associated with both GFAP+ and GFAP-cells, and were surrounded by a strong β-catenin expression (Fig. 4*E*, *middle panels, red*). We also observed fragmented laminin deposits in their interior (Fig. 4*E*, *arrowheads*). Serial imaging through the giant pinwheels and 3D reconstruction revealed that the giant bulbs associated with these clusters were filled with GFAP+ processes that transversed them and reached more distant and smaller fractones (Fig. 4*F*, *top and bottom panels*). These data indicate that fractones are not static structures, since their size, shape, position and cellular interactions change gradually with age, becoming larger and more complex in old mice.

**Figure 4:**
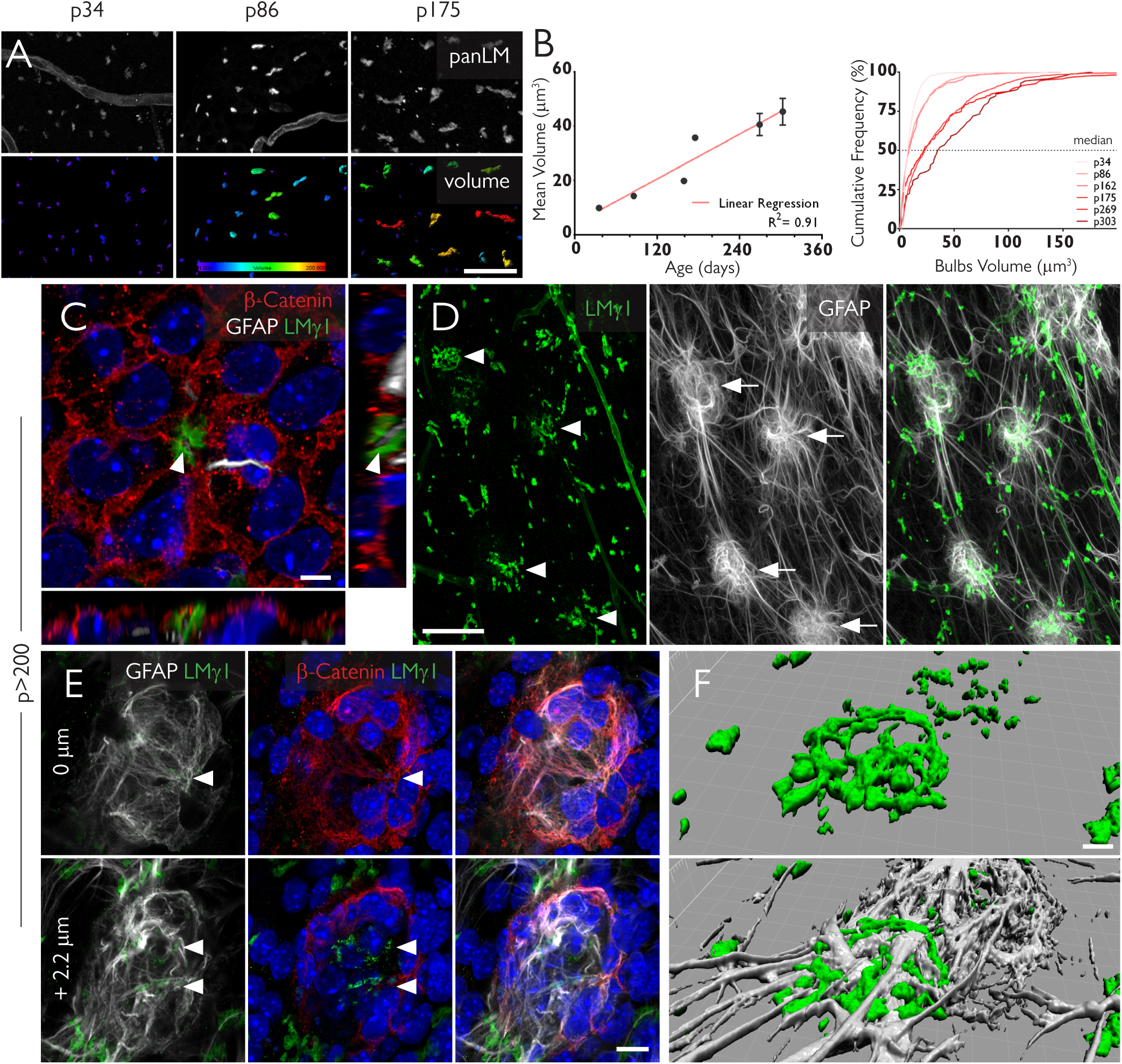
Fractones undergo major transformations with aging. ***A***, Fractones bulbs, identified by immunofluorescent staining using a pan-laminin antibody (panLM) (top panel), at three different ages. The volume of each individual bulb is color-coded, showing that these structures increase in size with age. ***B***, The mean volumes of fractones correlate with age (R²= 0.91, left panel). An analysis of frequency showing that the median value for the volume of fractones also increases with age. ***C***, A pinwheel in a whole mount of an 8 months old mouse showing a tunneled fractone bulb at its center. Orthogonal views show a GFAP+ process (gray) sheathed by the fractone BM (green). ***D***, Unusually large fractones (green, arrowheads) are associated with clusters of GFAP+ cells (gray, arrows). ***E***, A detailed view of a cell cluster shows strong expression of β-catenin (red) in its margins and fractones fragments (arrowheads) in its interior. ***F***, A tridimensional reconstruction of a giant fractone associated with a cell cluster showing how GFAP+ processes transverse these structures. Scale bars: A: 30 µm, C: 5 µm, D: 50 µm, E: 10 µm, F: 5 µm

The early appearance of laminin α5 in fractones bulbs, before its expression occurs at CNS blood vessels (Sorokin et al., 1997b) suggests an ependymal source of laminin α5+ fractones. To more precisely investigate this possibility, we investigated laminin α5+ fractone bulbs in transgenic mice lacking the expression of *Lama5* in endothelium (Tek-Cre::*Lama5*^−/−^) and in ciliated epithelial cells including ependymal cells (FoxJ1-Cre::*Lama5^−/−^*). Tek-Cre::*Lama5*^−/−^ mice showed no changes in laminin α5 staining of fractone bulbs (Fig. 5*A*). We quantified the volumes of bulbs in 3D reconstructions of confocal stacks and we did not find any significant differences among these genotypes (31.11µm^3^± 7.21 in WT vs. 29.22µm^3^ ± 16.1 in Tek-Cre::*Lama5*^−/−^) (Fig. 5*B*), suggesting that endothelial cells are not the cellular source of fractone bulbs. However, FoxJ1-Cre::*Lama5*^−/−^ showed almost complete loss of laminin α5 staining at fractone bulbs by 1 month of age, and only very few small laminin α5+ fractones (Fig. 5*C*, *left panel, arrowheads*). The activation of FoxJ1 promoter is temporally and spatially heterogeneous, starting at P0 and continuing until the third week after birth (Jacquet et al., 2009). Therefore, the few laminin α5-containing bulbs persisting in one-month old mutants were probably produced before the deletion was triggered by the FoxJ1 activation.We next investigated if laminin α5 containing bulbs were still present in FoxJ1-Cre::*Lama5*^−/−^ mice at two months of age. We found no laminin α5 in fractone bulbs, while it was still present in association with endothelial BMs (Fig. 5*D*, *center*).

**Figure 5:**
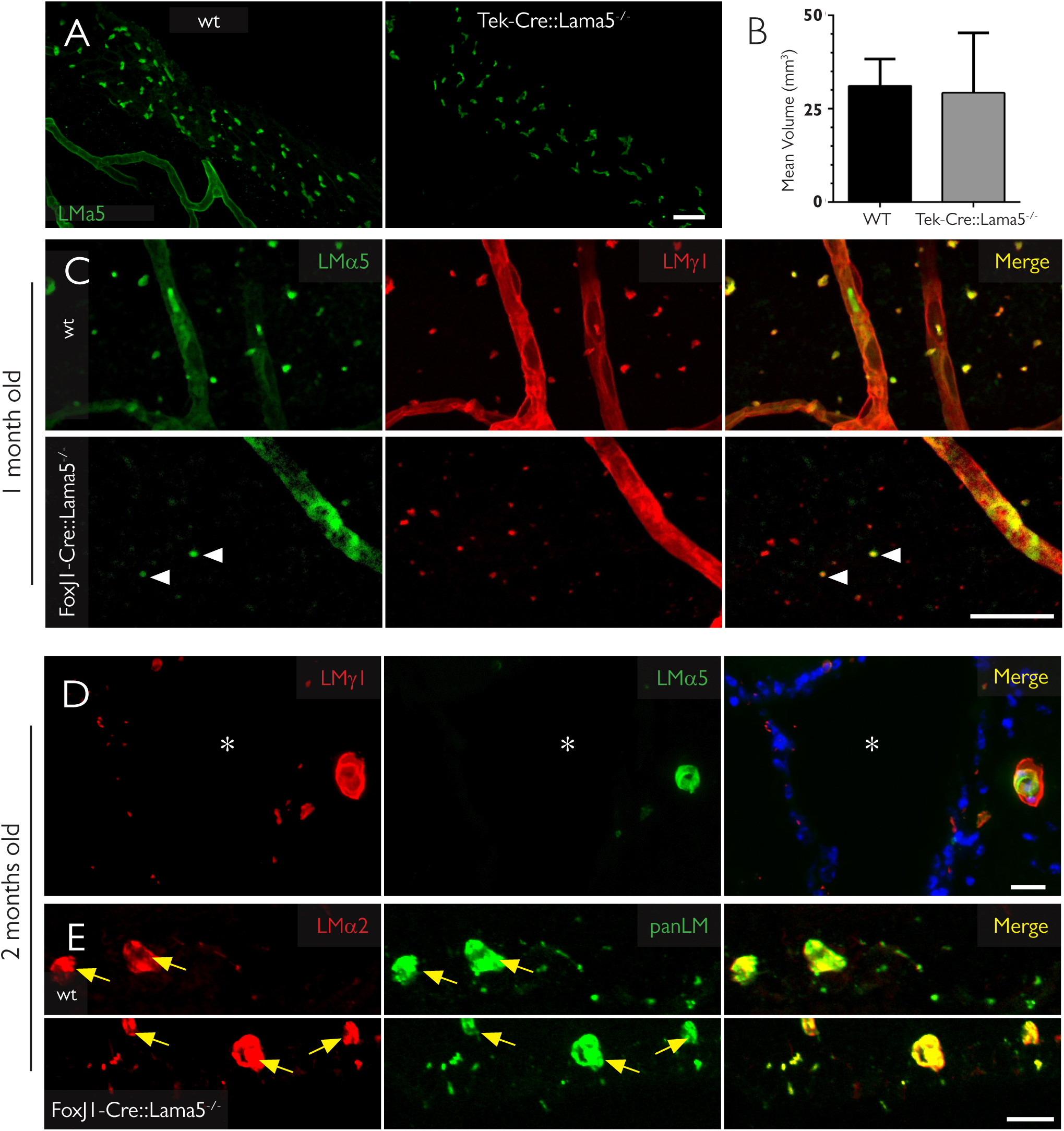
Deletion of the *Lama5* gene in ependymal cells eliminates laminin α5 in fractones bulbs. ***A***, Maximum intensity projection of 50 µm thick coronal sections of 5 months old mice showing that the expression of laminin α5 (green) is not altered in mice lacking the *Lama5* gene in endothelial cells (Tie2/*Lama5*^−/−^), indicating that these cells are not the source of the laminin α5 found in fractones. ***B***, Analysis of the volume of fractones showing no difference in bulb size among these genotypes (data are mean ± SEM, t-test n=3) ***C***, The vast majority of fractones in one month old FoxJ1/*Lama5*^−/−^ lack laminin α5, the main α chain in wild type animals. ***D***, Two month old FoxJ1/*Lama5*^−/−^ mice have no laminin α5 in fractone bulbs, instead, there is a compensatory upregulation of laminin α2 in these structures, (***E***) with a immunoreactivity similar to the blood capillaries in the vicinity (yellow arrows). Scale bars: A: 25 µm, C: 25 µm, D: 20 µm, E: 20µm.

Despite the lack of laminin α5 in FoxJ1-Cre::*Lama5*^−/−^ mice, fractone bulbs were still present as indicated by positive staining for other laminin chains, such as laminin γ1 (Fig 5*D*) and β1 chains and the other BM components such as collagen type IV and perlecan (not shown). Moreover, there were no detectable alterations in the general architecture of the ependymal layer, as judged by β-catenin immunoreactivity (not shown). Since laminins are heterotrimers, this finding suggests that other α chains could be expressed in fractones to compensate for the lack of laminin α5. Screening for the expression of the other laminin α chains, revealed upregulation of laminin α2 at fractones bulbs of FoxJ1-Cre::*Lama5*^−/−^ mice, which is almost absent at this site in wild-type animals (Fig 5*E*).

### FoxJ1-Cre: *Lama5*^−/−^ mice exhibit fewer slow-cycling NSCs and increased cell proliferation in the SVZ

As laminin α5 has been shown to maintain embryonic stem cells in a non-differentiated state and to regulate cell proliferation (Domogatskaya et al., 2008), we investigated whether the observed changes in laminin isoforms expression in the FoxJ1-Cre::*Lama5*^−/−^ mice could interfere with the physiology of the neural stem cell niche. We evaluated the number of NSCs in the SVZ by pulsing the thymidine analog, EdU, and tracking labelled cells after six weeks, using whole mounts of lateral wall preparations from WT and FoxJ1-Cre::*Lama5*^−/−^ mice (Fig. 6*A*). In this set up only slowly or non-dividing cells retain high levels of EdU. We found a significant 18% reduction in the number of NSCs in FoxJ1-Cre::*Lama5*^−/−^ compared to WT mice (4.95 +−0.20 × 10^−8^ cells/µm^2^ vs. 4.04 +−0.27 × 10^−8^ cells/µm^2^, p=0.02) (Fig. 6*B*). We also compared the total number of dividing cells in the SVZ, as assessed by phospho-histone H3 (PH3) staining, revealing a 60% increase in numbers of dividing cells when laminin α5 was absent from the fractone bulbs (3.42 +−0.77 × 10^−8^ cells/µm^2^ vs. 5.48 +−0.45 × 10^−8^ cells/µm^2^, p=0.04) (Fig. 6*C*). These results indicate that deletion of laminin α5 expression in ependymal cells reduces the quiescent stem cell pool, provoking an increase in proliferation of progenitor cells.

**Figure 6:**
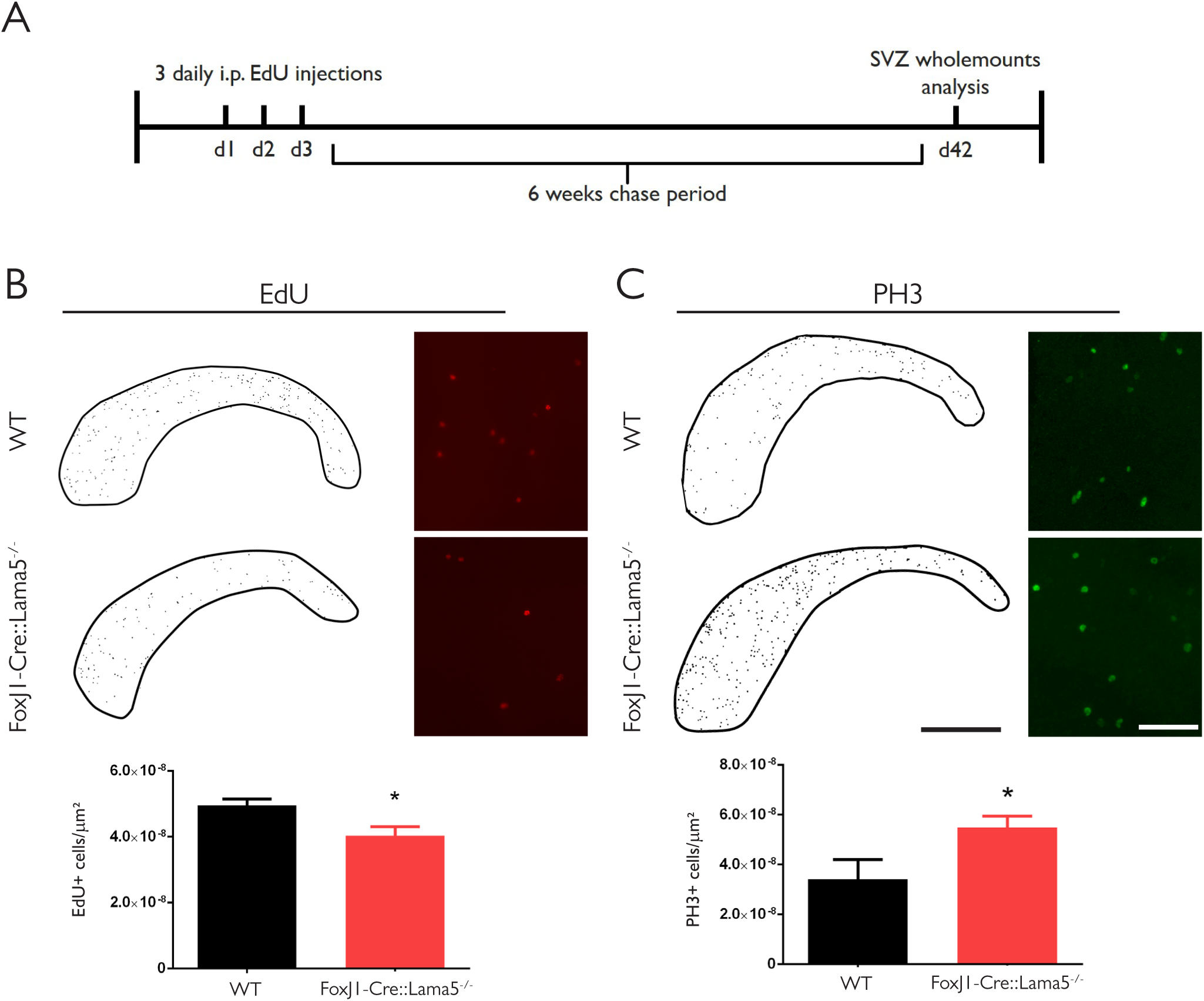
aminin content of fractone bulbs affect NSPC behavior in the SVZ. ***A***, EdU was administered to 3 months old mice for 3 days, and the 6 following weeks were a chase period. After that, SVZ wholemounts were analyzed in an epifluorescence microscope and Edu retaining cells and PH3+ cells were counted. ***B***, Counting of label retaining cells in the lateral ventricle of WT and FoxJ1/ *Lama5*^−/−^ revealed a reduction in the number of quiescent neural stem cells (from 4.95± 0.20x10^−8^ cells/µm^2^ N=6 to 4. 04 ± 0.27 ×10^−8^ cells/µm^2^ N=8; p=0.02). ***C***, Wholemounts were also immunofluorescently stained for PH3, a mitosis marker. FoxJ1/*Lama5*^−/−^ animals have a significant increase in the number of dividing cells comparing to wild type animals (5.48 ± 0.45x10^−8^ cells/µm^2^ N=5 vs. 3.42 ± 0.77 ×10^−8^ cells/µm^2^ N=5; p=0.04). N refers to individual wholemounts. Scale bars: B/C: low magnification: 1mm, high magnification: 100 µm.

## Discussion

We demonstrate here that a major laminin component of the fractone bulbs in the SVZ is laminin α5-containing isoforms, which is produced by the ependymal cells, and that the laminin content of fractones affects cell division in the SVZ. We additionally show that bulbs and stems, the two components of fractones, may represent independent structures, since they have distinct ECM compositions and are not always connected.

Fractones have been described as structures that emerge from the BMs of blood vessels in the vicinity of the SVZ. Given their expression of laminin α5, which is expressed in the CNS mainly by endothelial cells and deposited into the endothelial BM (Sorokin et al., 1997b; Sixt et al., 2001) it was logical to postulate endothelial cells as the source of fractones (Shen et al., 2008). However, in addition to endothelium, laminin α5 is one of the major BM components of epithelial cells (Sorokin et al., 1997a, 1997b) and has been previously described to be expressed by choroid plexus epithelium (Sorokin et al., 1997b). The use of transgenic mice lacking laminin α5 from either endothelial or ependymal cells revealed that the main source of laminin α5 and presumably also fractone bulbs are the ependymal cells lining the ventricle. We show here that fractone bulbs are in the ependymal cell layer, often in the center of pinwheels at the interface between neural stem cells and CSF. Our data also indicate that these interactions are dynamic, since fractone bulbs increase in size and their association with the GFAP+ neural stem cells increases in complexity with age.

Our data also showed that elimination of laminin α5 from fractone bulbs increased overall cell proliferation in the SVZ and reduced the number of slow dividing neural stem cells. After NSC activation and division its progeny leaves the SVZ within approximately 4–5 days (Ponti et al., 2013). Since the chase period used here was 6 weeks, cells retaining EdU in the SVZ corresponds to the small fraction of quiescent NSCs undergoing division within the 3-day period of the EdU pulse. Hence, those cells that were reduced in 18% are quiescent NSCs (inactive B cells), that were likely activated and started cycling faster in the absence of laminin α5. A possible explanation for the laminin α5-mediated maintenance of NSC quiescence could be through and indirect effect on TGF-β signaling. Although not clear how, it has been shown that the laminin α5 containing isoform laminin-511 strongly enhancesTGF-β signaling in epidermal stem cells (Morgner et al., 2015). TGF-β is an important factor that promotes quiescence in adult neural stem cells and its expression correlates with proliferation inhibition in the hippocampus (Kandasamy et al., 2014). The concomitant 60% increase in total cell proliferation in the SVZ in FoxJ1-Cre::*Lama5*^−/−^ is consistent with this hypothesis. However, we cannot exclude that lack of laminin α5 induces two separate effects, inhibiting activation of B cells and independently stimulating proliferation of neural progenitors. Nevertheless, our results indicate a role for laminin α5-containing fractone bulbs in the maintenance of the slow dividing neural stem cell pool.

Loss of laminin α5 correlated with a broader expression of laminin α2 in fractones. While the current data does not permit distinction between and anti-proliferative effect of laminin α5 or pro-proliferative effect of laminin α2, the former is consistent with data from other tissues suggesting that laminin α5 is a central factor in other stem cell niches. Stem cell niches in epithelia, bone marrow and the vascular niche in the hippocampus, are all rich in laminin α5 (Miner et al., 1997; Palmer et al., 2000; Kiel et al., 2005). Selective deletion of laminin α5 from keratinocytes has recently been shown to provoke hyperproliferation of cells in the basal layer of epidermis (Wegner et al., 2016). Laminin α5 containing isoforms also inhibit differentiation of embryonic stem cells, maintaining them in an anti-proliferative early progenitory state, through integrin signaling (Domogatskaya et al., 2008; Rodin et al., 2010; Hongisto et al., 2012). Consistent with an effect of laminin α5 on maintenance of neural stem cells quiescence, laminin receptors integrinα6β1, which has a higher affinity for laminin α5 compared to laminin α2 containing isoforms (Nishiuchi et al., 2006), and alpha-dystroglycan, are also associated with fractones (Shen et al., 2008; Adorjan and Kalman, 2009) and are downregulated on neural stem cells upon activation (Codega et al., 2014). Delivery of antibodies against this integrin into the CSF leads to a loss of NSC anchorage and increases its proliferation in 50% (Shen et al., 2008) and the proliferation of undifferentiated Sox2+ cells in 40% (Kazanis et al., 2010). However, it was assumed that the endothelial BM was the only source of ligands for the integrin α6β1, eventhough it is not clear whether stem cell processes directly contact laminin α5 in the endothelial BM.

As part of the blood-brain barrier, astrocytes ensheath capillaries in the CNS and produce a parenchymal BM rich in laminin α2, concealing the endothelial BM from other cells in the brain parenchyma (Sixt et al., 2001). It is not clear if neural stem cells, due their astrocytic phenotype, contribute to this parenchymal BM in the SVZ. In either scenario, the endothelial BM would not be fully available to integrin binding by NSC. In addition, we show here that stem cells can establish far more contacts with fractones than with blood vessels, hence, most of the exposed ligand for integrin α6β1 in NSC is the laminin α5 in fractone bulbs. We therefore propose that the integrin α6β1 positive neural stem cells interact with laminin α5 in fractones, which maintains neural stem cell character and limits proliferation and differentiation.

We also showed that pinwheels, formed by adjacent ependymal cells, can accommodate fractone bulbs at their centers where neural stem cell processes contact the CSF. We presume that these fractones provide an adherence spot for ependymal cell bodies and the apical processes of neural stem cells. In addition, the pinwheel center is a privileged site for tuning stem cell behavior, since the primary cilia at the tip of the apical process works as a signaling hub (Ihrie and Alvarez-Buylla, 2011). Having a BM at this critical site may help to stabilize the apical process in contact with the ventricle, exposing it to growth factors accumulated in fractones and enabling molecules at very low concentrations in the CSF to regulate neurogenesis. This interaction between apical processes and fractones increases in complexity during aging, as tunneled bulbs fully wrap the apical ending of stem cells. We also observed an increase in size of fractone bulbs with increased age of mice, confirming and extending recent findings which described a reduction in fractone bulb numbers with age (Kerever et al., 2015). The observed correlation between enlarged bulbs and GFAP+ cells clusters, which indicates that GFAP+ cells could participate in the merging of these structures, may result in the reduced fractone number reported by Kerever and co-workers. Taken together with our findings, which suggest an effect of laminin α5 in maintaining stem cell quiescence, the fusion of fractone bulbs could account for the age-related increase in neural stem cell activation, stem cell depletion and decline of neurogenesis (Shook et al., 2012).

In summary, our work shows that laminin α5-containing fractones are a critical element in the SVZ niche, being part of pinwheels and modulating the proliferation and stemness of NSPCs. Our data suggest that laminin α5 in fractones acts as a key stem cell niche factor. Further analyses of FoxJ1-Cre::Lama5−/−mice, investigating in detail the activation states of neural stem cells and changes in NSPC populations will aid in understanding the exact role of laminin α5 in fractones.

## Acknowledgements

Cells-in-Motion Cluster of Excellence (EXC1003) from the German Research Foundation (DFG). Brasilien-Zentrum of WWU/ DAAD for the financial support. National Council for Scientific and Technological Development (CNPq, 238147/2012-6). Dr. Eva Korpos for discussion, Dr Bhavin Shah for help with whole mounts and Stefan Luetke Enking for technical assistance. *FoxJ1-Cre* mice were a generous gift from Dr. Martin Bähler.

## Notes

**Conflict of Interest**: The authors declare no competing financial interests.

